# TFEB-depletion rescues TMEM106B-related myelin defects in zebrafish

**DOI:** 10.1101/2025.10.10.681547

**Authors:** W.M. Berdowski, H.C. van der Linde, N.I. Wolf, T.J. van Ham

## Abstract

Lysosomal transmembrane protein TMEM106B is genetically and neuropathologically implicated in a range of neurodegenerative diseases. Unexpectedly, the recurrent de novo TMEM106B variant p.(Asp252Asn) causes a hypomyelinating disorder. TMEM106B controls lysosomal positioning in neurons, but its role in white matter health is not fully understood. To study the role of TMEM106B in myelination in vivo, we mutated zebrafish *tmem106ba* and *tmem106bb* homologs. In vivo brain imaging in larvae deficient for both homologs (*tmem106b*^dm^) showed more, smaller lysosomes, accompanied by reduced myelin content and shorter myelin sheaths. Oligodendrocytes in *tmem106b*^dm^ exhibited more lysosomes in the perinuclear region. We next genome-edited the pathogenic p.(Asp252Asn) variant into the zebrafish genome and showed that it acts dominant-negatively on myelination and on lysosomal clustering around the nucleus in oligodendrocytes. As lysosomal peri-nuclear positioning modulates mTOR-signaling, and downstream transcription factor EB (TFEB) inhibits myelination, we next investigated whether there is a genetic interaction between TMEM106B and TFEB. Indeed, deficient myelination in *tmem106b^dm^* larvae, but not altered lysosomal clustering, was restored by loss of Tfeb in triple mutant larvae. Our work elucidates that TMEM106B modulates myelination by a TFEB-dependent pathway. This implicates the TFEB pathway in myelin development and white matter health and suggests that modulating TFEB could be a therapeutic target in white matter disorders.

## INRODUCTION

Lysosomal transmembrane protein 106B (TMEM106B) is an important player in neurodegenerative diseases. Genetic variants in TMEM106B modulate the risk for developing frontotemporal lobar degeneration TDP-43 pathology (FTLD-TDP), Alzheimer’s disease (AD), Parkinson’s disease (PD) and brain aging [392]. Recent studies further implicate TMEM106B in neurodegeneration, as amyloid fibrils from FTLD-TDP, AD and PD patients, but also of aged individuals with normal neurology, were primarily composed of TMEM106B filaments rather than TDP-43 [393–396]. TMEM106B is highly expressed in the central nervous system (CNS), including in neurons and oligodendrocytes, and localizes in the membrane of late-endosomal/lysosomal vesicles [104, 397, 398]. Evidence from in vitro and animal models shows that TMEM106B can regulate lysosomal biogenesis and lysosomal function, including morphology, enzyme activity, acidification, exocytosis, trafficking and positioning of lysosomes [102, 104–111]. The role of TMEM106B in neurodegeneration underscores the need to gain insight TMEM106B functions and general lysosomal dysfunction in the CNS and the relevance of this knowledge across multiple brain disorders.

Apart from its role in neurodegeneration, emerging evidence suggests a critical role for TMEM106B in myelinated axon bundles and white matter health. Recurrent *de novo* missense variant p.(Asp252Asn) in the luminal domain of TMEM106B causes hypomyelinating leukodystrophy (HLD) characterized by a significant lack of myelin and neurological symptoms and signs including congenital nystagmus and mild cognitive and motor delay [100, 399–401]. Myelin is formed by oligodendrocytes, which wrap their large membrane sheaths around axons in a multi-layered manner, thereby providing the basis for fast signal conduction and energy conservation, while simultaneously offering metabolic support and protection to axons [12]. *Tmem106b*-/-mice also show abnormal myelination and had lower levels of myelin proteins in brain lysate [107, 109, 402]. Evidence from a range of other genetic disorders affecting the white matter further implicates lysosomal regulation and function in white matter homeostasis, as mutations in various lysosomal enzymes and lysosomal fusion proteins can cause severe white matter diseases [67, 74, 75, 403]. However, it remains poorly understood how TMEM106B deficiency and lysosomal dysfunction lead to myelin abnormalities.

Defective lysosomal positioning and/or trafficking could play a role in the pathogenesis of TMEM106B-related HLD, but in vivo evidence is lacking. Myelin biogenesis requires lysosomal transport and exocytosis of myelin membrane proteolipid protein (PLP) [404–406]. The Asp252Asn (D252N) variant is located near the most distal glycosylation site (Asn256) of TMEM106B, which is particularly important for intracellular trafficking to late-endosomes/lysosomes [398]. Additionally, knock-down of TMEM106B and overexpression of TMEM106B containing the p.(Asp252Asn) variant caused lysosomal clustering in the perinuclear region of oligodendrocyte-like precursor cells in vitro [107, 109, 111]. Lysosomal positioning around the nucleus is associated with inactivity of mammalian target of rapamycin complex 1 (mTORC1), a multicomponent protein kinase that regulates many biosynthetic and catabolic processes [93, 407, 408]. mTORC1 is recruited by Rag-Ragulator complexes to lysosomes, and thereby can phosphorylate and regulate transcription factor EB (TFEB), a master transcriptional regulator of lysosomal biogenesis and autophagy [88–91, 409]. Both mTORC1 and TFEB modulate myelination. Signaling by the mTOR pathway promotes myelination, as activated mTOR causes excess myelin, while loss of mTOR signaling leads to hypomyelination [94–99]. In contrast, TFEB has a suppressive regulatory role in myelination, as TFEB deficient zebrafish and mice exhibit hypermyelination [85, 86]. Interestingly, TMEM106B may modulate TFEB localization and activity [102, 103]. Hence, we hypothesize that TMEM106B might modulate myelination via TFEB signaling.

Here, using zebrafish as a model system, we aimed to study how pathogenic *TMEM106B* variants affect lysosomal biogenesis and myelination in early CNS development, and whether TFEB signaling plays a role in TMEM106B-dependent myelin regulation. Zebrafish have been used extensively to study developmental genetics and mechanisms of myelination [52, 53, 280]. We generated *tmem106b* knock-out zebrafish and mutant zebrafish carrying the highly conserved p.(Asp252Asn) variant in *tmem106ba*. By real time in vivo imaging, we observed several oligodendrocyte-specific abnormalities in these mutants, including lack of myelin and lysosomal clustering around the nucleus, that might reveal novel lysosomal mechanistic pathways underlying TMEM106B-related hypomyelination. Intriguingly, loss of Tfeb could restore myelination, but not peri-nuclear lysosomal clustering in *tmem106b* knock-out fish. Together, these data raise a model wherein TMEM106B modulates myelination in oligodendrocytes through TFEB signaling, possibly by regulating lysosomal positioning.

## RESULTS

### Tmem106b deficiency causes more, smaller acidic vesicles in embryonic zebrafish brain

Two homologs of TMEM106B exist in zebrafish, Tmem106ba and Tmem106bb, sharing 62% and 72% sequence identity with the human protein respectively (Fig. 1A; Suppl. Fig. 1). Both homologs are expressed during embryonic development, although *tmem106ba* is highly expressed from a few hours post fertilization, while *tmem106bb* starts to be expressed at higher levels from 5 days post-fertilization (dpf) onwards (Fig. 1B) (NCBI BioProject No. PRJEB1986). As one homologous gene in zebrafish can compensate for loss of the other [410], we generated by CRISPR/Cas9 genome editing *tmem106b* double mutant (*tmem106b*^dm^) zebrafish, which contain a 44 bp insertion in *tmem106ba* and a 47 bp deletion in *tmem106bb* introducing a frameshift and premature stop (Fig. 1C, D). Since *tmem106ba* is highly expressed from early development, we studied developmental hallmarks in offspring of an incross of *tmem106ba^+/-^*zebrafish in a *tmem106bb*^-/-^ background. Overall developmental hallmarks, including body length, were normal in *tmem106b*^dm^ and in *tmem106ba^+/-^; bb^-/-^* at 5 dpf, indicating that Tmem106b deficiency does not cause major developmental delay, which is consistent with results obtained in *Tmem106b*-/-mice [107] (Suppl. Fig. 2A, B).

**Fig. 1.**
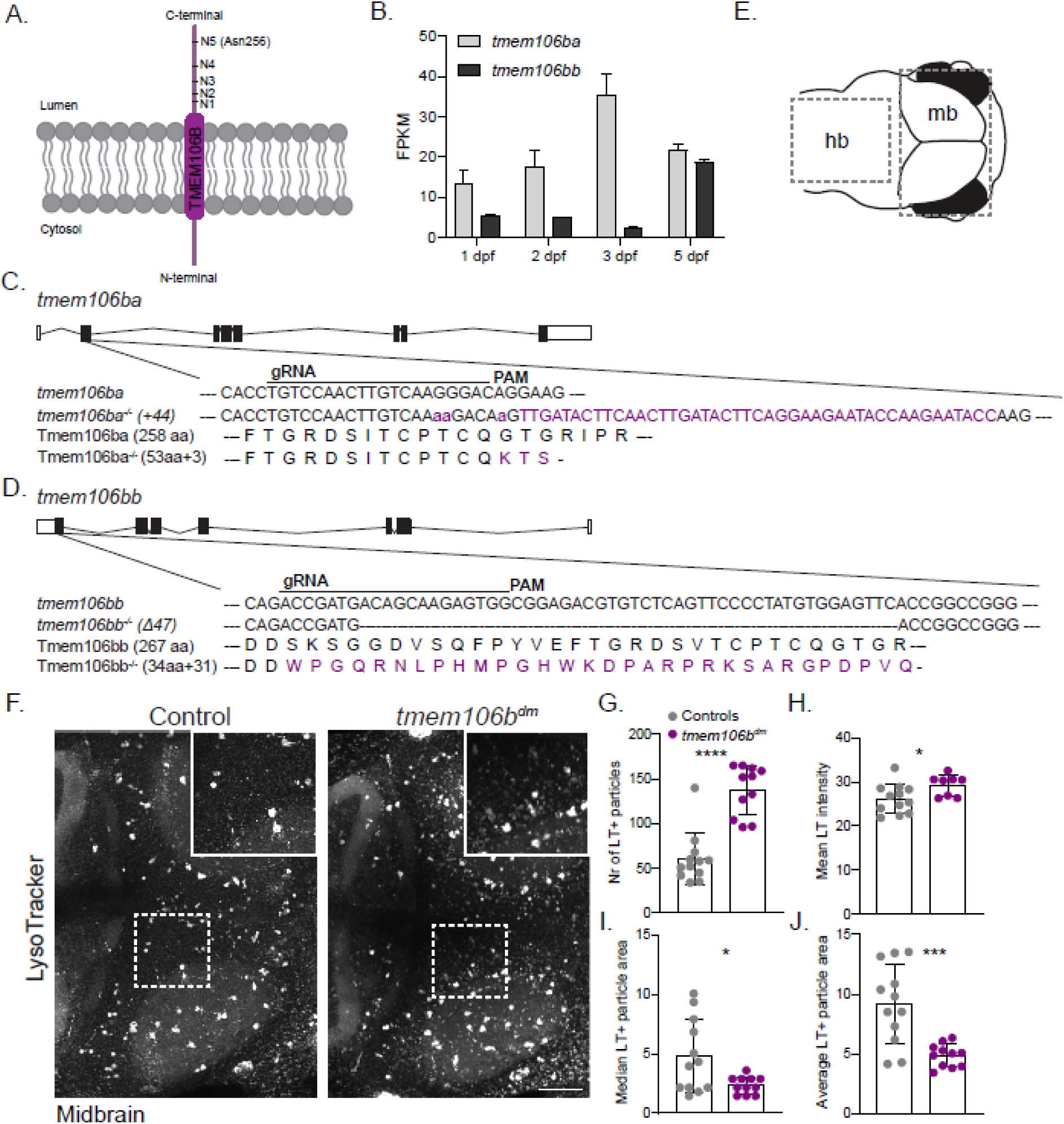
Loss of Tmem106b results in more and smaller LT+ particles in embryonic brain of zebrafish. A) Schematic of transmembrane protein TMEM106B, localized in the membrane of (endo-)lysosomes, including the five glycosylation sites in the luminal C-terminus. B) Expression levels of the two TMEM106B zebrafish homologs at 1 to 5 dpf, showing that *tmem106ba* is higher expressed during early embryonic development. C, D) Schematic representation of the two homologs of zebrafish *tmem106b*, the gRNAs (underlined) and PAM motif (magenta). *tmem106ba*^−/−^ fish have a 44 bp out of frame insertion in exon 2 (magenta DNA sequence), resulting in 53 + 3 out of frame amino acids (magenta protein sequence), followed by a premature stop codon (C). *tmem106bb*^−/−^ fish 47 bp out of frame deletion in exon 1, resulting in 34 + 31 out of frame amino acids (magenta DNA sequence) followed by a premature stop codon. E) Schematic representation of larval zebrafish head, indicating the brain regions imaged in this study: the hindbrain (hb) and midbrain (mb). F) Representative in vivo confocal images of LT staining in the midbrain of WT controls and *tmem106b*^dm^ larvae at 5 dpf. Insets show zoom-in of dashed square. Scale bar equals 50 µm. G, H, I, J) Quantification of the number of LT+ particles (G), average LT+ area (H), median LT+ area (I) and mean LT intensity (J) in the midbrain (n=10/group). Student’s t-test was preformed to test for significance (p < 0.05). hb: hindbrain; LT: LysoTracker; mb: midbrain. * p<0.05, *** p<0.001, **** p<0.0001.

We wondered whether loss of Tmem106b would cause lysosomal abnormalities in embryonic vertebrate brain development. We previously showed in multiple zebrafish lysosomal disease models that abnormal lysosomal phenotypes are most prominent in the midbrain [61, 73–75]. Therefore, we visualized acidic particles – including lysosomes – in vivo using LysoTracker Red (LT) and imaged the midbrain of controls (WT) and *tmem106b*^dm^ larvae at 5 dpf (Fig. 1E, F). *Tmem106b^dm^* larvae displayed increased numbers of LT+ particles and LT signal intensity in the midbrain compared to controls (Fig. 1G, H), indicating the presence of more lysosomes. These LT+ vesicles were predominantly located in between the two optic tecta—an area with a high density of radial glia, retinal axons and microglia—where they appeared to accumulate (inset Fig. 1F). Particle analysis in this region showed that *tmem106b*^dm^ larvae had more, but smaller LT+ particles compared to controls (Fig. 1I, J). Thus, Tmem106b deficiency results in more and smaller lysosomes in embryonic zebrafish brain, indicating that these mutants could serve as a relevant model to study TMEM106B function in early brain development.

### Tmem106b-deficiency causes fewer microglia and altered lysosomes in microglia

Microglia are the major phagocytes of the brain and - in particular in brain development - they are packed with lysosomes and exhibit high lysosomal activity to assist in degradation of phagocytic debris and apoptotic cells. Microglia first colonize the zebrafish midbrain around 2 dpf and can be visualized by transgenic *mpeg1*:GFP and Neutral Red (NR) staining [181, 323]. To study putative microglial abnormalities, we first quantified NR+ and transgenically labeled *mpeg1:*GFP + microglia in the midbrain of controls and *tmem106b*^dm^ larvae. We found that *tmem106b*^dm^ larvae had fewer NR+ and *mpeg*:GFP+ microglia in the midbrain compared to controls at 3 and 5 dpf (Fig. 2A, B; Suppl. Fig. 2C, D). To limit potential effects of experimental or genetic background differences, and because *tmem106ba* is highly expressed around the initiation of microglial colonization, we then incrossed *tmem106ba^+/-^*fish in a *tmem106bb* knock-out background and quantified NR+ microglia in their progeny. Heterozygous mutant *tmem106ba* larvae had normal microglial numbers at 5 dpf, whereas *tmem106b*^dm^ fish showed fewer microglia compared to controls and heterozygotes (Fig. 2C, D). As expected, progeny of a *tmem106ba^-/-^; bb^+/-^* incross does not show altered microglial numbers, indicating that *tmem106ba* could be more important in these early developmental stages (Suppl. Fig. 2E, F).

**Fig. 2.**
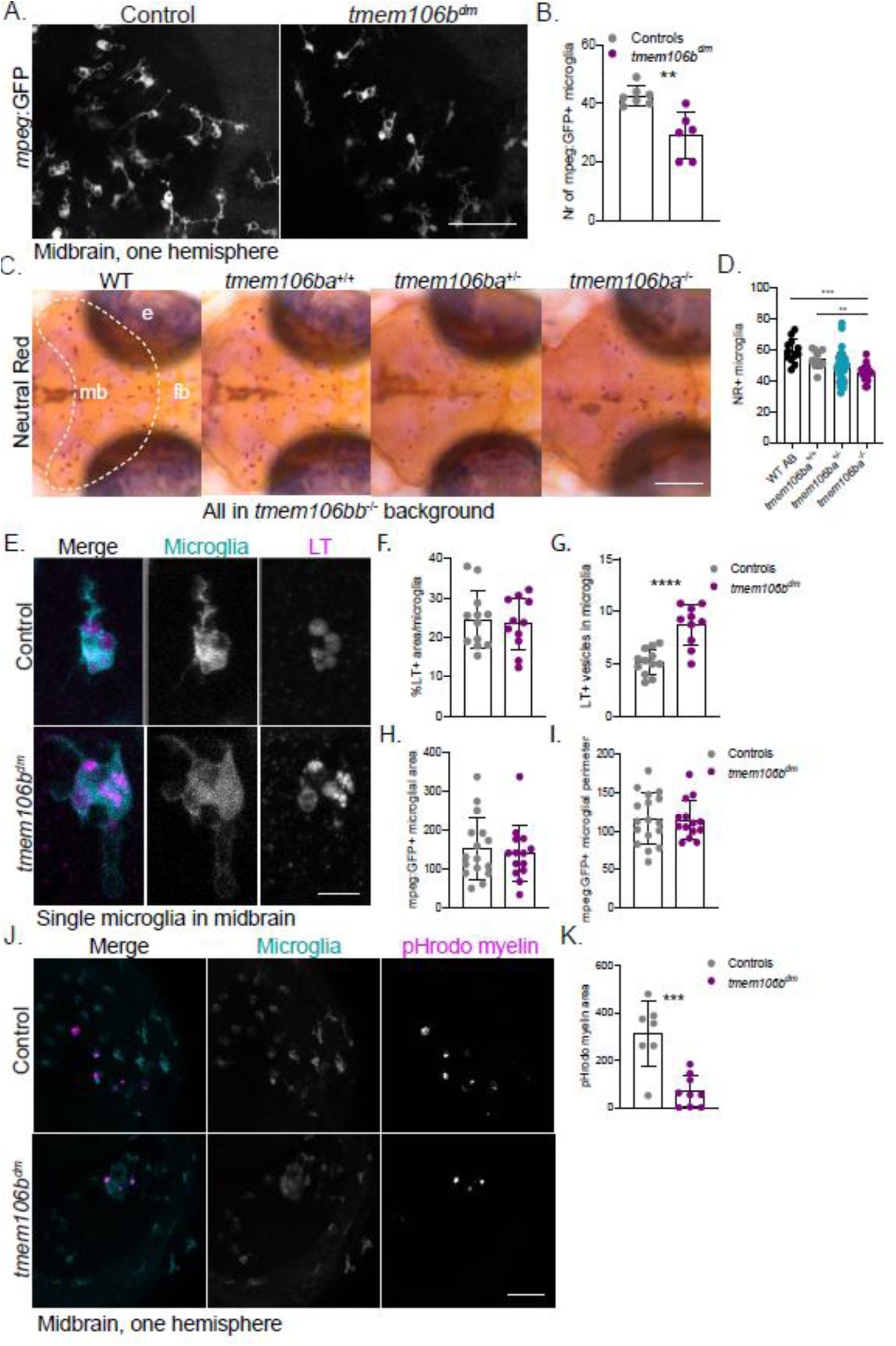
Loss of Tmem106b causes loss of microglia and altered lysosomes in microglia. A) Representative in vivo confocal images of *mpeg1*:GFP + microglia in one hemisphere of the midbrain of controls and *tmem106b*^dm^ larvae at 5 dpf. Scale bar equals 50 µm. B) Quantification of *mpeg*:GFP+ microglia in controls (n = 7) and *tmem106b*^dm^ (n = 6) larvae. C) Representative images of the midbrain (mb, dashed line) after NR staining of WT AB larvae, and offspring of *tmem106ba*^+/-^ *tmem106bb*^-/-^ incross, at 5 dpf. Scale bar equals 200 µm. D) Quantification of NR+ microglia in WT AB (n = 12), *tmem106ba*^+/+^ *tmem106bb*^-/-^ (n = 10), *tmem106ba*^+/-^ *tmem106bb*^-/-^ (n = 23) and *tmem106b*^dm^ (n = 16) larvae at 5 dpf. E) Representative in vivo confocal images of LT staining (magenta) of WT and *tmem106b*^dm^ larvae in a Tg(*mpeg*:GFP) background, showing single microglia (cyan) in the midbrain at 5 dpf. Scale bar equals 10 µm. F, G) Quantification of the % LT+ area in microglia (F) and the number of LT+ vesicles in microglia (G). H, I) Quantification of area (H) and perimeter (I) of *mpeg*:GFP+ microglia. J) Representative in vivo confocal images of *mpeg1*:GFP + microglia (green) in the midbrain after intracerebral injection (6 hpi) with pHrodo-labelled myelin (magenta) of controls and *tmem106b*^dm^ larvae at 3 dpf. Scale bar equals 50 µm. K) Quantification of pHrodo-labelled myelin area in microglia. One-way ANOVA test with multiple comparisons and Student’s t-test were preformed to test for significance (p < 0.05). e: eye; fb: forebrain; LT: LysoTracker; mb: midbrain. ** p<0.01, **** p<0.0001.

Next, to study putative lysosomal abnormalities in microglia, we stained transgenic *mpeg1*:GFP zebrafish of 5 dpf in control and *tmem106b*^dm^ background with LT (Fig. 2E). Although LT area in *tmem106b*^dm^ microglia did not differ from controls, Tmem106b-deficient microglia contained more LT+ vesicles (Fig. 2E, F, G). To test whether these lysosomal alterations impact microglial function, we analyzed the morphology of *mpeg1*:GFP+ microglia in the midbrain at 5 dpf. A more round, amoeboid microglial morphology indicates increased phagocytic activity and is associated with brain pathology [2, 411]. In normal conditions, Tmem106b-deficient microglia showed a normal ramified morphology, as total cellular area and perimeter were not different from controls (Fig. 2H, I). However, microglia in *Tmem106b*-/- mice showed impaired proliferation and activation in response to demyelination [412]. To study whether microglia in *tmem106b*^dm^ larvae respond differently to myelin debris, we performed intracerebral injections of pHrodo-labelled human myelin in transgenic *mpeg1*:GFP controls and *tmem106b*^dm^ larvae and quantified the uptake of myelin in microglial phagosomes. Six hours post-injection, microglia in *tmem106b*^dm^ larvae had less pHrodo-labelled myelin area in microglia, suggesting a decline in myelin debris uptake and lysosomal degradation (Fig. 2J, K). Altogether, loss of Tmem106b causes a reduction in microglial expansion, and more small LT+ vesicles in microglia, without prominent morphological in normal condition, but reduced ability to phagocytose myelin debris.

### Loss of Tmem106b affects earliest myelination and lysosomal clustering in oligodendrocytes

We next wondered whether *tmem106b*^dm^ display altered myelination phenotypes associated with hypomyelination. We crossed *tmem106b*^dm^ fish with transgenic *mbp*:eGFP-CAAX fish, in which membrane bound GFP is driven by the myelin basic protein (MBP) promoter, to visualize myelinating oligodendrocytes [293]. We performed in vivo imaging and observed that *tmem106b*^dm^ larvae had decreased *mbp*:GFP+ myelin area in the hindbrain at 5 dpf, which was accompanied by shorter and fewer myelinated axons perpendicular to the two Mauthner axons (Fig. 3A, B, C, D; Mauthner axons indicated with M). Interestingly, *tmem106ba^+/+^; bb^-/-^* and *tmem106ba^+/-^; bb^-/-^* larvae both had more myelin content in the hindbrain than their *tmem106b*^dm^ siblings, showing that heterozygous loss of Tmem106b is insufficient to cause hypomyelination (Suppl. Fig. 2G, H). To independently assess myelin development, we used whole mount *in situ* hybridization (WISH) for *mbpa*, which showed less staining consistent with hypomyelination in *tmem106b*^dm^ larvae (Fig. 3E). WISH for *plp1b*, which is translated in oligodendrocytic somata and can therefore be used to visualize myelinating oligodendrocytes, showed normal staining, indicating that hypomyelination in *tmem106b*^dm^ larvae is independent of oligodendrocyte loss (Fig. 3F). Since TMEM106B is known to play a role in axonal trafficking, the decreased myelin content in the hindbrain could be secondary to abnormal axonal development or loss of axons [104, 108]. We therefore performed whole-mount immunohistochemistry for acetylated tubulin on larvae to visualize axons, and imaged the midbrain optic tectum, which has a large density of axons. We found no indication of major axonal pathology or axon loss in *tmem106b*^dm^ larvae at 5 dpf, which implies that the reduced myelination is not caused by widespread axonal loss (Fig. 3F). Altogether, this suggests that *tmem106b*^dm^ larvae have a lack of myelin independent of loss of oligodendrocytes or axons.

**Fig. 3.**
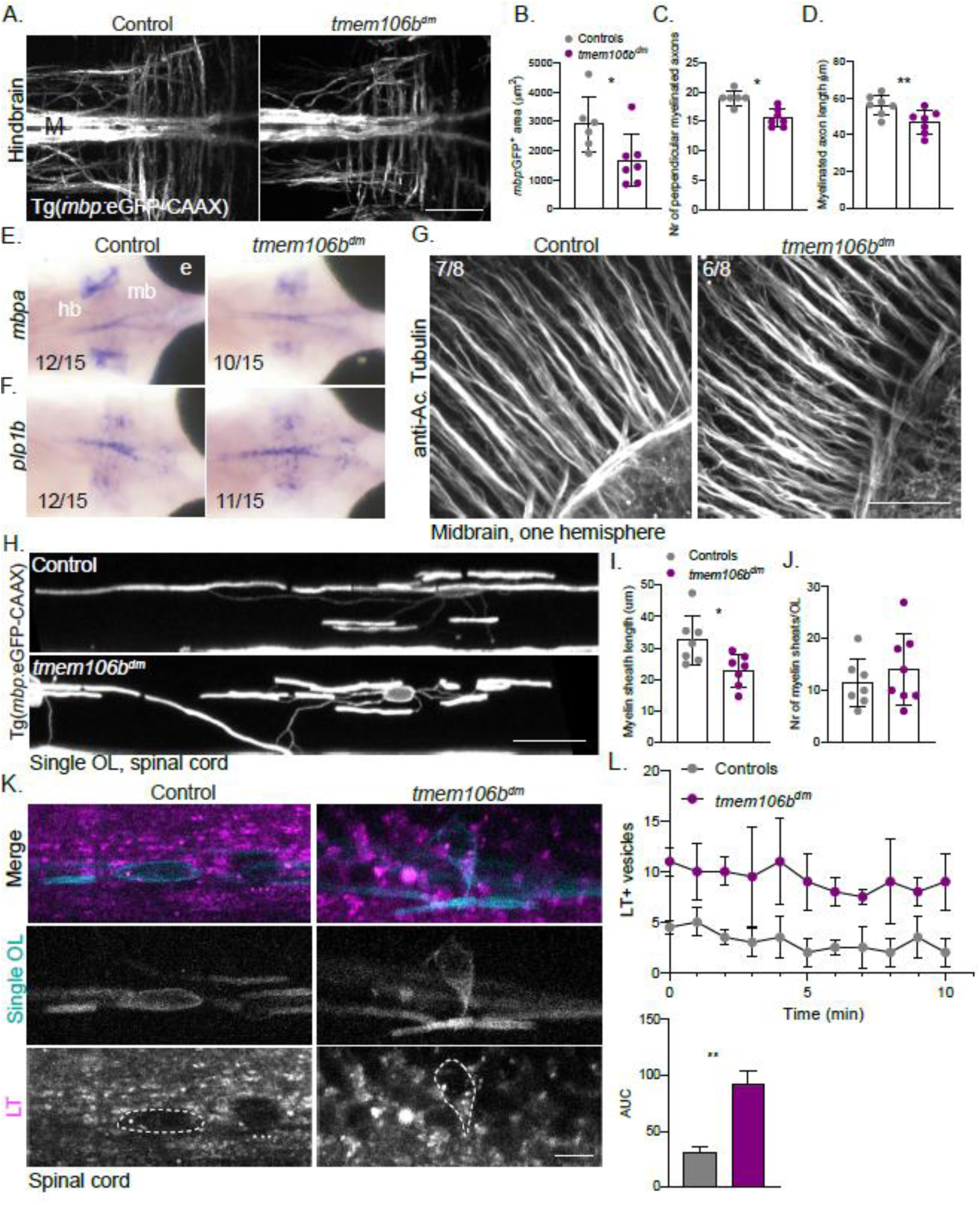
Loss of Tmem106b causes lysosomal clustering in oligodendrocytes and hypomyelination independent of loss of oligodendrocytes or axons. A) Representative in vivo confocal images of *mbp*:GFP+ myelin in the hindbrain of controls and *tmem106b*^dm^ larvae at 5 dpf. Scale bar equals 50 µm. B, C, D) Quantification of mbp:GFP+ area (B), number of perpendicular myelinated axons (C) and myelinated axon length (D) in controls (n = 6) and *tmem106b*^dm^ (n = 7) larvae at 5 dpf. E, F) Representative images of in situ hybridization of *mbpa* (E) and *plp1b* (F) in brain of controls and *tmem106b*^dm^ larvae at 5 dpf, showing reduced *mbpa* + myelin sheaths (10/15 larvae) but similar number of mature *plp1b*+ oligodendrocytes in *tmem106b*^dm^ larvae (11/15 larvae). G) Representative confocal images of the midbrain (one hemisphere) of controls and *tmem106b*^dm^ larvae (n = 8/group) stained with anti-Acetylated Tubulin, to visualize axons, showing no substantial axonal loss in *tmem106b*^dm^ larvae (6/8 larvae). Scale bar equals 20 µm. H) Representative in vivo confocal images of individual *mbp*:GFP+ myelinating oligodendrocytes in the spinal cord of control and *tmem106b*^dm^ larvae at 5 dpf, showing shorter myelin sheaths in Tmem106b deficient oligodendrocytes. Scale bar equals 20 µm. I, J) Quantification of length (I) and the number (J) of *mbp*:GFP+ myelin sheaths in control and *tmem106b*^dm^ larvae (n = 7/group) at 5 dpf. K) Representative in vivo confocal images of individual *mbp*:GFP+ myelinating oligodendrocytes (cyan) in the spinal cord of controls and *tmem106b*^dm^ larvae at 3 dpf, stained with LT (magenta). Images are obtained from a 10-minute timelapse video (Suppl. Video 1), and show altered positioning of lysosomes in the soma (dashed lines) of Tmem106b deficient oligodendrocytes. Scale bar equals 5 µm. L) Quantification of the number of LT+ particles in individual oligodendrocytes per minute (n = 2/group, top), over the course of a 10 minute timelapse video, and the area under the curve (AUC, bottom). Student’s t-test was preformed to test for significance (p < 0.05). hb: hindbrain; LT: LysoTracker; mb: midbrain. * p<0.05, ** p<0.01.

To study myelination in more detail, we imaged individual oligodendrocytes in the posterior spinal cord and measured individual myelin sheaths, as described before [293]. Tmem106b-deficient oligodendrocytes displayed shorter myelin sheaths compared to controls at 5 dpf, while the number of sheaths per cell did not differ (Fig. 3I, J). Loss of TMEM106B in primary oligodendrocytes, neurons and other cell lines in vitro leads to clustering of lysosomes around the nucleus [104, 107, 111]. To test whether this also happens in vivo, we performed in vivo time-lapse imaging of individual oligodendrocytes of control and *tmem106b*^dm^ larvae stained with LT at 3 dpf, a time point where spinal cord myelination starts in zebrafish [52]. Intriguingly, Tmem106b-deficient oligodendrocytes displayed clustering of LT+ vesicles in the perinuclear region, in particular where the soma membrane extends towards the myelin sheath (Fig. 3K). Particle analysis within the soma showed an increase in LT+ vesicle numbers in Tmem106b-deficient oligodendrocytes over time, based on the area under the curve (AUC) (Fig. 3L). The number of LT+ vesicles remained constant over ten minutes in control and *tmem106b*^dm^ oligodendrocytes (Slopes: Controls: -0,227, *tmem106b*^dm^: -0,255). Altogether, *tmem106b*^dm^ larvae showed shortened myelin sheaths, which appeared independent of loss of oligodendrocytes or axons, and more lysosomes in the perinuclear region in oligodendrocytes during early myelination.

### Missense variant p.(Asp252Asn) causes hypomyelination and lysosomal clustering in a dominant negative manner

Having shown that *tmem106b* loss of function causes several lysosomal and myelination abnormalities in zebrafish larvae, we further studied the recurrent de novo *TMEM106B* missense variant p.(Asp252Asn), which causes HLD [100]. Asp252 is located in the luminal domain of TMEM106B and is highly conserved among vertebrates, including zebrafish (Fig. 4A, B). To study early pathological mechanisms of the pathogenic Asp252Asn (D252N) variant, we used a CRISPR/Cas9 knock-in strategy and introduced p.(Asp252Asn) in the *tmem106ba* genomic locus in zebrafish with a *tmem106bb*^-/-^ background (Fig. 4C). Siblings of *tmem106ba*^D252N/+^ crossed with *tmem106ba*^ex1/+^ showed normal developmental hallmarks, including body length (Suppl. Fig. 3 A, B). First, we investigated myelination in *tmem106ba*^D252N/+^ larvae by crossing *tmem106ba*^D252N/+^ with *tmem106ba*^ex1/+^ mutants in Tg(*mbp*:eGFP-CAAX) background (Fig. 4D). As we showed before, *mbp*:GFP+ myelinated area in *tmem106ba*^ex1/+^ larvae does not differ from WT controls (Suppl. Fig. 1G, H). In contrast, *tmem106ba*^D252N/+^ larvae already exhibited significantly decreased myelinated area (Fig. 4D). *Tmem106ba*^D252N/+^ and *tmem106ba*^D252N/ex1^ larvae showed similar myelin content in the hindbrain (Fig. 4D). This points towards a dominant negative effect of the p.(Asp252Asn) variant on myelination, which is consistent with the finding that TMEM106B is capable of forming homo- and heterodimers [102].

**Fig. 4.**
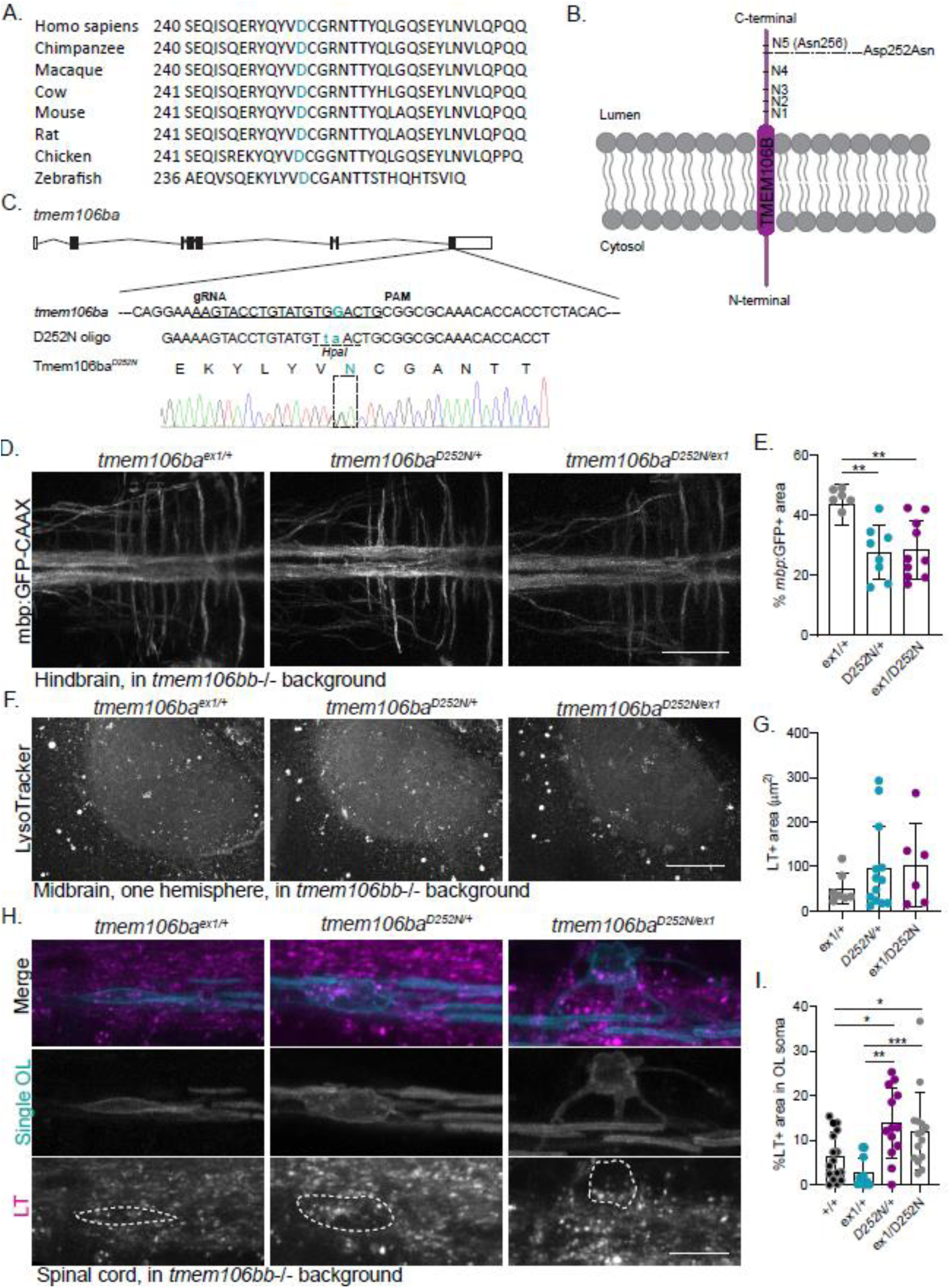
Missense variant p.(Asp252Asn) acts dominant negatively on myelin and lysosomal clustering in oligodendrocytes, but does not affect lysosomes in the brain. A) The mutated Asp residue is highly conserved across species, including in zebrafish. B) Schematic of TMEM106B, showing the location of the *de novo* missense variant Asp252Asn which causes hypomyelinating leukodystrophy in patients. C) Schematic representation of *tmem106ba* with: the location of the Asp252Asn missense variant located in exon 8, the 19 bp gRNA, the PAM motif and the co-injected 30 bp oligo containing the missense variant. This results in an Asp to Asn change at residue 252 (green) and a G>T at 251 (green) which generates a restriction site (HpaI) without changing the amino acid (Val>Val). Tmem106ba missense mutants are generated in a *tmem106bb*-/-background to prevent genetic compensation. D) Representative in vivo confocal images of *mbp*:GFP+ myelin in the hindbrain of *tmem106ba*^ex1/+^, *tmem106ba*^D252N/+^ and *tmem106ba*^D252N/ex1^ larvae at 5 dpf. Scale bar equals 50 µm. E) Quantification of the %mpb:GFP+ area in *tmem106ba*^ex1/+^ (n = 6), *tmem106ba*^D252N/+^ (n = 8) and *tmem106ba*^D252N/ex1^ (n = 10) larvae at 5 dpf. F) Representative in vivo confocal images of LT staining in the midbrain (one hemisphere) of *tmem106ba*^ex1/+^, *tmem106ba*^D252N/+^ and *tmem106ba*^D252N/ex1^ larvae at 5 dpf. Scale bar equals 50 µm. G) Quantification of LT+ area in one hemisphere of the midbrain *tmem106ba*^ex1/+^ (n = 7), *tmem106ba*^D252N/+^ (n = 12) and *tmem106ba*^D252N/ex1^ (n = 6) larvae at 5 dpf. H) Representative in vivo confocal images of individual *mbp*:GFP+ myelinating oligodendrocytes (cyan) in the spinal cord of *tmem106ba*^ex1/+^, *tmem106ba*^D252N/+^ and *tmem106ba*^D252N/ex1^ larvae at 3 dpf, stained with LT (magenta). Scale bar equals 10 µm. I) Quantification of the % LT+ area in the soma of individual oligodendrocytes, in *tmem106ba*^+/+^ (n = 16 OLs, 8 fish), *tmem106ba*^ex1/+^ (n = 11 OLs, 6 fish), *tmem106ba*^D252N/+^ (n = 13 OLs, 6 fish) and *tmem106ba*^D252N/ex1^ (n = 15 OLs, 8 fish) larvae at 3 dpf. One-way ANOVA test with multiple comparisons was preformed to test for significance (p < 0.05). * p<0.05, ** p<0.01, *** p<0.001. LT: LysoTracker; OL: oligodendrocyte.

To study whether p.(Asp252Asn) affects lysosomes, we performed LT staining of progeny of *tmem106ba*^D252N/+^ crossed with *tmem106ba*^ex1/+^ mutants. In contrast to *tmem106b*^dm^ larvae, we found no evidence for major lysosomal abnormalities in the midbrain of *tmem106ba*^ex1/+^, *tmem106ba*^D252N/+^ and *tmem106ba*^D252N/ex1^ mutants at 5 dpf (Fig. 4F, G). Contrarily, *tmem106ba*^D252N/+^ and *tmem106ba*^D252N/ex1^ larvae, but not *tmem106ba*^ex1/+^ larvae, showed increased LT+ area and altered lysosomal clustering around the nucleus in oligodendrocytes during the initial phase of myelination (Fig. H, I). Since *tmem106b*^dm^ mutants had fewer microglia, we quantified the number of NR+ microglia in 3 dpf offspring of *tmem106ba*^D252N/+^ crossed with *tmem106ba*^ex1/+^ mutants (Suppl. Fig. 3C). Curiously, microglia numbers did not differ between genotypes, in contrast to the decrease we observed in *tmem106b*^dm^ (Suppl. Fig. 3C, D). Altogether, our data suggest that the p.(Asp252Asn) variant specifically affects myelination and peri-nuclear lysosomal clustering in oligodendrocytes in a dominant-negative manner.

### Loss of Tfeb restores myelination, but not lysosomal clustering, in tmem106b^dm^ larvae

Having shown that oligodendrocytes in *tmem106b*^dm^ and *tmem106ba*^D252N/+^ had more lysosomes clustering around the nucleus, we further tested our hypothesis that TMEM106B could modulate myelination via the TFEB pathway. Loss of TFEB was previously shown to result in increased myelination [85, 86]. Indeed, we observed more *cldnk*:GFP+ myelin content in the hindbrain of *tfeb*^-/-^ larvae compared to controls, already at 5 dpf [85] (Fig. 5A, B). TMEM106B may modulate TFEB localization and activity [102, 103]. We hypothesized that TMEM106B deficiency might lead to activation of TFEB and thereby inhibit myelination (Fig. 5C). To explore whether there is a genetic interaction between TMEM106B and TFEB in myelination, we performed epistasis experiments using *tfeb*-/-mutants to test whether loss of Tfeb modulates the myelination phenotype in *tmem106b* mutants [85]. Intriguingly, *mbp*:GFP+ myelinated area in the hindbrain of *tmem106b*^dm^;*tfeb*^-/-^ larvae was increased compared to *tmem106b*^dm^ mutants, but did not differ from control animals, suggesting that loss of *tfeb* can restore myelination (Fig. 5D, E). To exclude possible experimental or genetic background effects, we compared *mbp*:GFP+ myelinated area in progeny of incrossed *tmem106ba*^+/-^;*bb^-/-^*; *tfeb*^+/-^ fish at 5 dpf, and observed that *tmem106ba*^+/-^;*bb^-/-^*; *tfeb*^+/-^ and *tmem106b*^dm^;*tfeb*^-/-^ larvae had more myelin content in the hindbrain compared to their *tmem106b*^dm^ siblings (Suppl. Fig. 4A, B). The myelinated area in *tmem106b*^dm^;*tfeb*^+/-^ mutants was not significantly different from either *tmem106b*^dm^ or *tmem106b*^dm^;*tfeb*^-/-^ siblings, suggesting a possible partial rescue (Suppl. Fig. 4A, B). Altogether, this shows that loss of Tfeb can restore myelination in *tmem106b*^dm^ larvae.

**Fig. 5.**
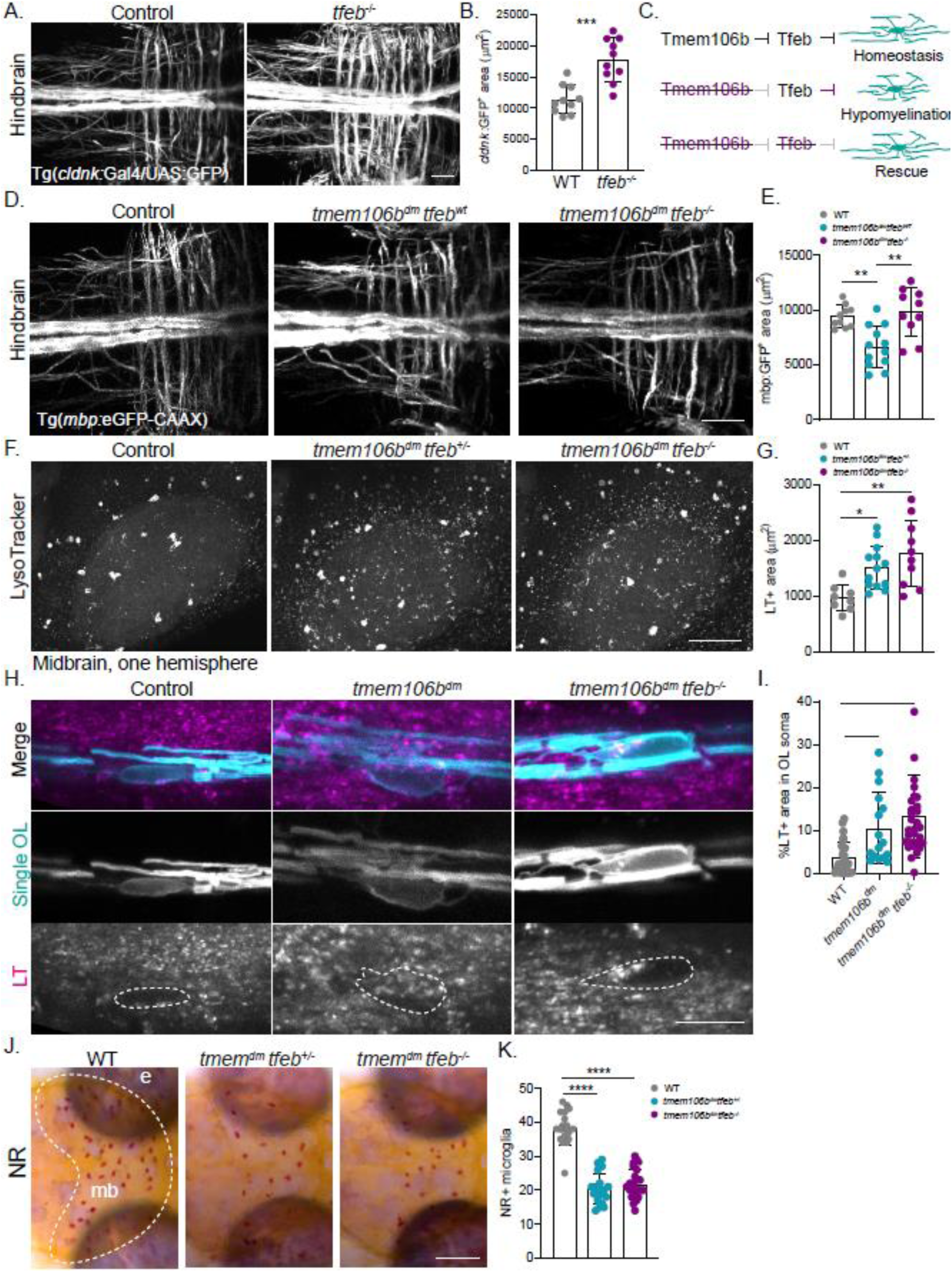
Loss of Tfeb restores myelination in *tmem106b*^dm^ larvae, but not lysosomal phenotypes. A) Representative in vivo confocal images of *cldnk*:GFP+ myelin in the hindbrain of controls and *tfeb*^-/-^ larvae at 5 dpf. Scale bar equals 20 µm. B) Quantification of the *cldnk*:GFP+ myelin area in the hindbrain of controls and *tfeb*^-/-^ larvae at 5 dpf (n = 10/group). C) Schematic representation of our hypothesis that Tmem106b regulates myelination by suppressing Tfeb. Loss of Tmem106b could lead to overactive Tfeb, which suppresses myelination. If this is true, loss of Tfeb in *tmem106b*^dm^ larvae should restore the hypomyelination phenotype. D, E) Representative in vivo confocal images (D) and mbp:GFP+ area quantification (E) of *mbp*:GFP+ myelin in the hindbrain of WT controls (n = 10), *tmem106b*^dm^ (n = 12) and *tmem106b*^dm^ *tfeb*^-/-^ (n = 10) larvae at 5 dpf. Scale bar equals 30 µm. F, G) Representative in vivo confocal images (F) and LT+ area quantification (G) of LT staining in the midbrain (one hemisphere) of WT controls (n = 8), *tmem106b*^dm^ *tfeb*^+/-^ (n = 13) and *tmem106b*^dm^ *tfeb*^-/-^ (n = 10) larvae at 5 dpf. Scale bar equals 50 µm. H, I) Representative in vivo confocal images of individual *mbp*:GFP+ myelinating oligodendrocytes (cyan) stained with LT (magenta) in the spinal cord (H) and quantification of the % LT+ area in oligodendrocyte soma (I) of WT controls (n = 26 OLs, 12 fish), *tmem106b*^dm^ (n = 16 OLs, 10 fish) and *tmem106b*^dm^ *tfeb*^-/-^ (n = 28 OLs, 12 fish) larvae at 3 dpf. Scale bar equals 20 µm. J, K) Representative images of the midbrain (mb, dashed line) after NR staining (J) and NR+ microglia quantification (K) of WT larvae (n = 19), *tmem106b*^dm^ *tfeb*^+/-^ (n = 20) and *tmem106b*^dm^ *tfeb*^-/-^ (n = 23) at 3 dpf. Scale bar equals 200 µm. One-way ANOVA test with multiple comparisons or Student’s t-test was preformed to test for significance (p < 0.05). * p<0.05, ** p<0.01, *** p<0.0001, **** p<0.0001.

We then wondered whether Tfeb deficiency also rescues the lysosomal abnormalities in *tmem106b*^dm^ mutants. Similar to the observation in oligodendrocytes derived from *Tfeb*-/-mice, loss of Tfeb did not affect the number or morphology of LT+ vesicles in the zebrafish midbrain at 5 dpf (Suppl. Fig 4. C, D) [86]. We compared the LT+ area in the midbrain of controls and progeny of *tmem106b*^dm^;*tfeb*^+/-^ fish crossed with *tmem106b*^dm^;*tfeb^-/-^*, and found that the lysosomal phenotype is not restored in *tmem106b*^dm^*tfeb^-/-^* larvae (Fig. 5F, G). Equally, oligodendrocytes in *tmem106b*^dm^;*tfeb^-/-^* fish showed increased percentage LT+ area in their soma and perinuclear clustering, similar to what we observed in *tmem106b*^dm^ larvae (Fig. 5H, I). Altogether, this implies that in *tmem106b*^dm^, loss of Tfeb may restore myelination downstream of lysosomal clustering in oligodendrocytes.

Microglia numbers are diminished in zebrafish mutant for RagA, one of several GTPases that can recruit mTORC1 to the lysosome surface, and this phenotype is partially rescued by knock-out of *tfeb* [413]. We therefore quantified NR+ microglia in controls and offspring of *tmem106b*^dm^ *;tfeb^-/-^* crossed with *tmem106b*^dm^*;tfeb^+/-^*, and found that loss of Tfeb did not rescue the decrease in microglia numbers, nor did it rescue impairment of pHrodo-labelled myelin phagocytosis (Fig. 5J, K; Suppl. Fig. 4 C, D). This indicates that loss of Tfeb might cell-specifically restore myelin biogenesis in Tmem106b-deficient zebrafish.

## DISCUSSION

Here, we established genetic zebrafish models to explore TMEM106B function and the role of pathogenic missense variant p.(Asp252Asn), causing a hypomyelinating disorder in humans, on early myelination and lysosomal biogenesis in early brain development. Our data indicate that loss of Tmem106b in vivo causes lysosomal alterations, including increased numbers of lysosomes localized around the nucleus in myelinating oligodendrocytes. Moreover, we showed that Tmem106b depletion causes reduced myelination and shortened myelin sheath development, and that the p.(Asp252Asn) missense variant acts dominantly in myelinating oligodendrocytes possibly in a cell-specific manner. Mechanistically, we provide in vivo evidence that there is a genetic interaction between TMEM106B and key lysosomal regulator TFEB, as TMEM106B-related hypomyelination can be restored by depleting Tfeb. This pathogenic cascade could be initiated by increased lysosomal numbers around the nucleus in oligodendrocytes and consequent activation of lysosomal transcription factor TFEB [93]. Suppressing TFEB-signaling may provide a route to counteract myelination deficiency due to TMEM106B mutations or possibly other pathogenic mechanisms involving lysosomes and myelination.

We demonstrated that *tmem106b* mutant zebrafish can serve as a relevant model to study TMEM106B function in early brain development, as they had more, smaller lysosomes in the brain, and lysosomal clustering around the nucleus in oligodendrocytes, which is consistent with observations in previous in vitro models [102, 107, 109, 111, 414]. Mutant *tmem106b* larvae showed hypomyelination in the CNS accompanied with shorted myelin sheaths, that appeared to be independent of loss of oligodendrocytes or axons. *Tmem106b*-/-mice also exhibited hypomyelination, although other reported normal myelination [109, 402, 415]. Additionally, *Tmem106b*-/-mice had reduced expression of genes encoding myelin sheath proteins, but not of Olig2, a transcription factor which is highly expressed in the whole oligodendrocyte lineage, indicating that indeed TMEM106B deficiency particularly affects myelin sheaths, independent of oligodendrocyte loss [107, 402]. Moreover, axon pathology is profound in aged *Tmem106b*-/-mice, but not in 2-month-old mice, indicating that this is an age-related phenotype [402, 416]. Hence, we find it unlikely that the hypomyelination in Tmem106b deficient larvae is caused by primary axonal loss. However, we cannot exclude involvement of axonal abnormalities, as TMEM106B depletion in neurons in vitro causes several neuronal abnormalities including reduced dendrite branching and defective axonal lysosomal trafficking [102, 104, 108]. Altogether, loss of TMEM106B appears to particularly affect myelin sheath development.

Our data revealed a genetic interaction between TMEM106B and lysosomal transcription factor TFEB in regulating myelination, as loss of Tfeb restores myelination in *tmem106b*^dm^ larvae. TFEB is predominantly expressed in pre-myelinating oligodendrocytes and mature oligodendrocytes [86], and Tfeb depletion does not restore microglial numbers. Hence, these restorative effects appear cell autonomous to oligodendrocytes. Intriguingly, Tfeb depletion also does not rescue altered lysosomal clustering in oligodendrocytes. As perinuclear localization of lysosomes is associated with mTORC1 inactivity, perinuclear clustering of lysosomes might precede Tfeb activation in Tmem106b-deficient oligodendrocytes, which in turn could suppress myelination [85, 86, 93]. However, previous studies showed that several other kinases can also phosphorylate TFEB [90, 417–421], including AKT, which is also inhibited by perinuclear clustering of lysosomes, and highly involved in myelination [96–98, 408]. Nonetheless, circumstantial evidence does support a link between TMEM106B and mTORC1 activation. TMEM106B interacts with vacuolar H+– adenosine triphosphatase ATPase (v-ATPase) via accessory protein 1 (AP1), which is necessary for amino acids to activate mTORC1 through Rag-Ragulator recruitment of mTORC1 to the lysosomal surface [106, 422]. Of note, the p.(Asp252Asn) variants does not appear to affect interaction with AP1, but it is unclear whether it affects v-ATPase functioning [109]. Altogether, we provided in vivo evidence that Tfeb depletion can restore myelination in Tmem106b-deficient zebrafish, but how TMEM106B precisely modulates the TFEB pathway remains to be established. It would be interesting to chemically or photogenetically manipulate the localization of lysosomes in oligodendrocytes, to further explore how lysosomal positioning affects TFEB signaling and myelination [423–425].

There are several possible mechanisms how enhanced TFEB activity could induce hypomyelination. TFEB is a master regulator of autophagy and promotes endocytosis [88, 89, 409, 426]. Oligodendrocytic endocytosis of their own myelin sheaths, and subsequent autophagy and myelin turnover, are all required for proper myelin development, myelin plasticity during motor learning and for maintaining healthy myelin throughout adult life [86, 188, 379, 427–429]. Interestingly, TMEM106B deficiency is also associated with increased autophagy and impaired maturation of autophagosomes to autolysosomes [108, 110, 111, 415, 430]. Moreover, TFEB regulates expression of several lysosomal hydrolases, which are crucial for lysosomal recycling in autophagy [88]. Tmem106b depletion in mice causes increased protein levels of cathepsin D and cathepsin B, and elevated enzyme activity of Hexa and Glb1, which are all TFEB target genes [108–110, 402]. Hence, TFEB might suppress myelination in TMEM106B-related hypermyelination by enhancing autophagic lysosomal degradation pathways. Besides promoting autophagy and endocytosis, TFEB is known to promote apoptosis of pre-myelinating oligodendrocytes [86]. However, we find it unlikely that enhanced apoptosis underlies hypomyelination in *tmem106b* mutant zebrafish, as we found no indication for oligodendrocyte cell death. It will be intriguing to further explore how TFEB regulates myelination, both in TMEM106B-related hypomyelination and in normal brain development.

We suspected that the Asp252Asn variant acts dominant negatively, as TMEM106B can form homo- and hetero dimers with TMEM106C [102]. Additionally, TMEM106B-related HLD patient fibroblasts and cells overexpressing TMEM106B^D252N^ had normal RNA and protein levels, indicating that the p.(Asp252Asn) variant does not affect RNA or protein stability [109, 431]. Indeed, we demonstrated in vivo that the heterozygous p.(Asp252Asn) missense variant, but not a heterozygous null allele, results in hypomyelination and peri-nuclear lysosomal clustering in oligodendrocytes. The p.(Asp252Asn) variant, and the nearby fifth C-terminal glycosylation site (Ans256), appears to significantly affect lysosomal positioning and trafficking [109, 398]. However, it remains unknown how exactly the variant impacts TMEM106B functioning. Curiously, unlike *tmem106b^dm^*larvae, the heterozygous p.(Asp252Asn) variant failed to induce altered lysosomal morphology or numbers in the midbrain, in line with previous in vitro studies [109]. Also, heterozygous p.(Asp252Asn) mutants did not show less microglia than controls, unlike *tmem106b^dm^.* This strongly suggests that the Asp252Asn variant acts dominant-negatively on myelin formation and the amount of lysosomes surrounding the nucleus in oligodendrocytes in particular.

Myelination requires coordinated lysosomal trafficking, uptake, storage and release of myelin proteins, including PLP, which may be affected by altered lysosomal clustering in TMEM106B deficient oligodendrocytes [404, 406, 432]. Indeed, TMEM106B deficiency results in less PLP on the lysosomal surface, indicating inhibited lysosomal exocytosis, while overexpression of TMEM106B enhances secretion of lysosomal proteins [103, 109]. Curiously, Tfeb depletion in *tmem106b* mutants did not affect altered lysosomal clustering, but did restore myelination, which raises the question whether impaired lysosomal localization and/or trafficking is indeed detrimental to myelin development. Again, it would be interesting to manipulate lysosomal positioning in oligodendrocytes to study the effects on protein trafficking and myelination.

Tmem106b depletion in zebrafish diminishes microglia and affects microglial phagocytosis of myelin debris in response to increased phagocytic demand. The latter is consistent with recent observations that microglia in *Tmem106b*-/- mice showed impaired proliferation and activation in response to demyelination [412]. Aside from this, *Tmem106b*-/- mice displayed conflicting microglial phenotypes, as some report no differences in microglial numbers or proteins, while others noted more microglia and microgliosis in the brain [109, 110, 415, 430]. Interestingly, post-mortem corpus collosum tissue from patients carrying a TMEM106B risk allele also showed reduced microglial numbers [412]. We recently showed that loss of microglia causes an adult-onset leukodystrophy [59, 61]. Therefore, we cannot exclude possible effects of loss of microglia on myelination in *tmem106b*^dm^ larvae. However, the p.(Asp252Asn) missense variant did not affect microglia numbers in zebrafish larvae and it is yet unknown if TMEM106B-related HLD patients present with microglial abnormalities. Additionally, depletion of Tfeb could not rescue the loss of microglia in *tmem106b^dm^* zebrafish larvae, while it did restore myelination. Altogether, even though TMEM106B depletion may reduce microglial expansion and phagocytic capacities of microglia, this indicating that the p.(Asp252Asn) variant and the restorative effects of Tfeb depletion predominantly affect oligodendrocyte biology.

Zebrafish allow for detailed exploration of cells in embryonic development, in their natural environment by non-invasive in vivo examination. This has led to novel fundamental insights in myelination [31, 43, 44, 48, 49, 147, 268, 269, 433]. Possible limitations of zebrafish when studying white matter disorders include the lack of a layered cortex and a low ratio of myelinated versus non-myelinated brain tissue. Nevertheless, we were able to detect subtle myelin changes in our *tmem106b* mutant zebrafish that could be more easy to miss in other models. For example, overall myelination appears to be largely normal in the corpus callosum of 8-month-old *Tmem106b-/-* mice, while expression of myelinating oligodendrocyte-specific genes was already downregulated, which does already indicate subtle myelin defects [107]. Hence, as we show here, zebrafish in particular are suitable to detect phenotypes that go unnoticed in other models.

Concluding, we provide further mechanistic evidence for the role of TMEM106B in myelination, where TMEM106B may act by regulating localization and/or trafficking of lysosomes, and show that a pathogenic TMEM106B variant acts dominant negatively on the number of perinuclear lysosomes and myelination. Our data reveal a novel genetic link between TMEM106B and the TFEB pathway in regulating myelination, which may provide a novel route to modulate myelination in disease.

## MATRIALS & METHODS

### Zebrafish models

Zebrafish were maintained under standard conditions [322]. Adult fish were fed brine shrimp twice a day and kept in groups on a 14-h light and 10-h dark cycle. Until 5 dpf, zebrafish embryos and larvae were kept at 28 °C on a 14–10-h light–dark cycle in 1 M HEPES buffered (pH 7.2) E3 medium (34.8 g NaCl, 1.6 g KCl, 5.8 g CaCl2·2H2O, 9.78 g MgCl2·6H2O). To prevent pigmentation, the medium was changed to E3 medium containing 0.003% 1-phenyl 2-thiourea (PTU) (from now on referred to as E3/PTU) before 24 h post-fertilization (hpf).

The following mutant zebrafish lines were used: wild-type AB; wild-type TL; *tmem106b^dm^*with a 44 bp out-of-frame insertion in exon 1 of *tmem106ba* (*tmem106ba^+44/+44^*) and a 47 bp out-of-frame deletion in *tmem106bb (tmem106bb*^Δ47/Δ47^); *tmem106ba^D252N^* carrying the Asp252Asn substitution in exon 8 of *tmem106ba*, in a *tmem106bb*^Δ47/Δ47^ background to avoid genetic compensation.

The *tmem106b^dm^*were crossed with Tg(*mpeg1*:eGFP) zebrafish, expressing GFP under the control of the mpeg1 promotor to visualize microglia [323], and with Tg(*mbp*:EGFP-CAAX) expressing membrane-bound GFP under the control of the mbp promotor to visualize mature oligodendrocytes and myelin [293].

The *tfeb*^-/-^ mutants (*tfeb*^st120^) have an out-of-frame 5 bp deletion in exon 3, resulting in a premature stop (45aa+16) [85]. To analyze myelin content in these fish, *tfeb* mutants in a transgenic Tg(*cldnk*:Gal4/UAS:GFP-CAAX) background were used [85, 387]. Mutants without a transgenic background were used to cross in with *tmem106b^dm^* in a Tg(*mbp*:EGFP-CAAX) background, to generate *tmem106b^dm^ tfeb*^+/-^ and *tmem106b*^dm^ *tfeb*^-/-^ fish. Animal keeping was approved by the Animal Experimentation Committee at Erasmus MC, Rotterdam.

### CRISPR-Cas9 genome editing in zebrafish – Generation of tmem106b^dm^

CRISPR/Cas9 mutagenesis was performed using *tmem106ba* and *tmem106bb* specific guideRNA’s (gRNAs) targeting exon 2, that were designed and synthesized as previously described [186, 434] (gRNA *tmem106ba* target sequence: TGTCCAACTTGTCAAGGGACAGG; gRNA *tmem106bb* target sequence: ACCGATGACAGCAAGAGTGGCGG). The SP-Cas9 plasmid (Addgene plasmid #62731) was used for the production of Cas9 protein and was kindly provided by Niels Geijsen [338]. Cas9 nuclease was synthetized as described previously [338]. Cas9 nuclease was then co-injected with one of the gRNAs into fertilized zebrafish oocytes. Indel efficiency and frequency were determined using Sanger sequencing. Injected embryos were raised up and crossed out against wild-type (WT) fish. Founder fish were genotyped using PCR and Sanger sequencing, and selected for breeding. F1 fish with a 44 bp insertion in *tmem106ba* (pG54K fs 2*) or a 47 bp deletion in *tmem106bb* (S35W fs 30*) were then crossed to obtain stable mutant lines.

### CRISPR-Cas9 genome editing in zebrafish – Generation of Asp252Asn missense mutants

To introduce the highly conserved Asp252Asn missense variant causing HLD into *tmem106ba* in the zebrafish genome, we used CRISPR/Cas9 with co-injection of an ssDNA oligo [Integrated DNA Technologies (IDT), standard desalted oligo (STD)], as described previously [61, 337]. We used the Alt-R™ CRISPR-Cas9 System of IDT to generate a gRNA with target sequence AAGTACCTGTATGTGGACTG. Equal amounts of crRNA (crispr RNA) and tracrRNA (trans-activating crispr RNA) in Duplex Buffer (IDT) were incubated for 5 min at 95 °C, followed by cooling to RT to generate 50 µM gRNA duplex. The ssDNA oligo (GAAAAGTACCTGTATGTtaACTGCGGCGCAAACACCACCT) contained: 20 bp homology arms up- and downstream of the double-stranded break created by Cas9; the Asp252Asn missense variant (G>A) causing HLD; a G>T mutation (Val251Val) to generate a restriction site (HpaI) and prevent recutting of Cas9. The restriction site was used for genotyping.

For injections, 50 pmol gRNA was mixed with 4 ng Cas9 protein to form gRNA-Cas9 RNPs. Next, 30 pmol oligo and 0.4 µl of 0.5% Phenol Red was added in a total volume of 6 µl. One nl was injected in the zebrafish embryo at the one-cell stage. Indel efficiency was determined using Sanger sequencing. Injected embryos were raised up and crossed out against WT fish. F1 zebrafish with the Asp252Asn mutation in *tmem106ba* were crossed to obtain a stable mutant line with a *tmem106bb* knock-out background.

### Genotyping of zebrafish and zebrafish larvae

Adult zebrafish were anesthetized with 0.016% MS-222 and a small piece of the caudal fin was cut and lysed in 80 µl 50 mM KOH. Zebrafish larvae were euthanized and lysed in single tubes containing 80 µl 50 mM KOH per larva, followed by an incubation for 30 min at 95 °C. Eight µl 1 M Tris–HCl pH 8 was added, and 1 µl of the lysate was used for PCR (Touch-down PCR 65-55: 5′ 95 °C, 30″ 95 °C, from 65 °C to 55 °C in 10 cycles, 45″ 72 °C, 30″ 93 °C, 30″ 58 °C, 45″ 72°C repeated 25 times, 3′ 72 °C). For genotyping of *tmem106ba*^+44/+44^ and *tmem106bb*^Δ47/Δ47^, 5 µl of PCR product with 10 µl GelRed/OrangeG was loaded on a 2% Tris-borate-EDTA (TBE) agarose gel to separate the WT and mutant allele based on size (Table 1). For genotyping of *tmem106ba^D252N^*, we designed an allele specific PCR that required two PCR reactions. Both reactions contained a forward primer and reverse primer covering part of exon 8 with the Asp252Asn missense mutation. In one reaction, a forward primer for the mutant allele was added, and in the other reaction a forward primer for the WT allele. Five µl of product of each PCR reaction was loaded on a 2.2% TBE agarose gel (Table 1). See Table 1 for the primer sequences.

**Table 1.**
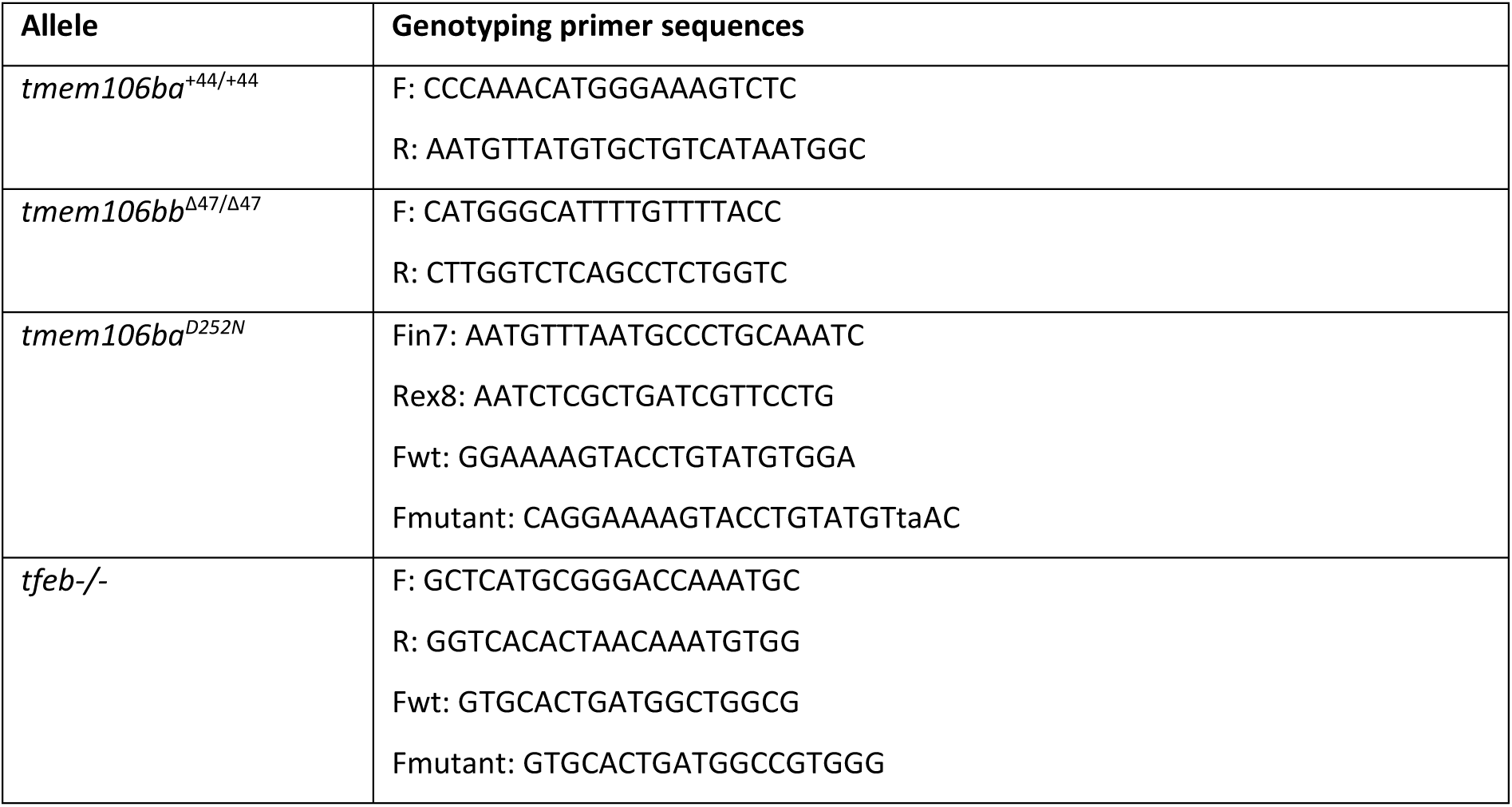
Primer sequences for genotyping

### LysoTracker Red staining

Live zebrafish larvae (n = ∼ 15) were incubated in 2 mL tubes with 10 μM LysoTracker™ Red DND-99 (1:100) (ThermoFisher, Waltham, MA) in 200 μl E3/PTU. Tubes with open lids were kept in the dark at 28 °C for 40 min. Larvae were then washed with E3/PTU for 10–15 min at 28 °C in the dark. We started in vivo imaging within 45 min after staining.

### In vivo confocal imaging acquisition

Live zebrafish larvae were anesthetized using 0.016% MS-222 and mounted in 1.8% low-melting-point agarose in E3 medium. During imaging, the dish was covered with E3 medium containing MS-222. We acquired all confocal images using a Leica SP5 intravital microscope with a 20 × water dipping objective (Leica Plan-Apochromat, NA = 1.0) using 488 nm, 561 nm and 633 nm lasers.

For myelin area quantification in the brain, larvae were embedded dorsally and z-stack images (∼70 z-stacks, step size 2 µm) were acquired in the hindbrain. For detailed analysis of single oligodendrocytes, larvae were embedded laterally and z-stack images (∼30 z-stacks, step size 0.5 µm) were acquired in the posterior spinal cord, where individual oligodendrocytes were well distinguishable. For in vivo timelapse imaging of LT+ vesicles in single oligodendrocytes, we acquired z-stack images (3 z-stacks, step size 0.5 µm) every 4 seconds for 10 minutes of single oligodendrocytes in the spinal cord at 3 dpf, a timepoint where myelination is initiated in zebrafish [52]. For analysis on LT+ vesicles in the brain and in microglia, images of dorsally embedded larvae were acquired (∼50 z-stacks, step size 2 µm) in the midbrain. If progeny from heterozygous crosses were used, larvae were sacrificed and genotyped after imaging as described in ‘Genotyping of zebrafish and zebrafish larvae’.

### Intra-cerebral pHrodo-labelled myelin injection

pHrodo-labelled myelin was injected intra-cerebrally in 3 dpf larvae in a Tg(*mpeg*:GFP) background as previously described [61]. Six hours post-injection (hpi), we obtained in vivo confocal images to assess the amount of pHrodo-labelled myelin that had been taken up by microglia. Red pHrodo-labeled myelin was kindly gifted by Inge Huitinga (Amsterdam UMC, The Netherlands Brain Bank). Briefly, myelin was isolated from post-mortem brain tissue of a pool of healthy control donors, and labelled with the pH-sensitive fluorescent dye pHrodo, as previously described [44].

### In situ hybridization

Whole-mount in situ hybridization to visualize *mbpa* and *plp1b* mRNA was performed as previously described [61, 339]. As described previously, riboprobes were prepared via T7 transcription using digoxigenin-labeled NTPs (Roche, Mannheim, Germany) and a cDNA template [340]. The primers used are listed below (underlined region indicates the T7 promoter sequence).

*mbpa*: F CTAAGTCGAGGGGAGAAAGCC / R: TAATACGACTCACTATAGGGAGGGCATACAATCCAAGCCA (product size 885 bp).

*plp1b*: F TCCTCTATGGACTGTTGCTGCTG / R TAATACGACTCACTATAGGGACAATCACACACAGGAGGACCAA (product size 1052 bp).

### Whole-mount acetylated tubulin staining and image acquisition

Five dpf-old larvae were fixed in 4% PFA at room temperature (RT) for 3 h, washed in PBS-T (1% Triton-X100 in PBS), dehydrated to 100% MeOH, and kept at -20°C for at least two hours. After rehydration to PBS-T, larvae were washed 4 x 5 min and permeabilized in ice cold acetone (prechilled at -20° freezer) for 5-8 min. After extensive washing, blocking buffer (1% BSA 0.1% DMSO in PBST) was added for 2 hr at RT. Afterwards, larvae were incubated with 5% BSA/PBST containing primary anti-body (1:500 mouse-anti α-tubulin α4a, Sigma, T6793) for 2 days at -4°C. After extensive washing with PBS-T (10 x 30 min), larvae were incubated with anti-mouse DyLight Alexa 647 (1:200) for 1-2 days. Hoechst was used for nuclear counter staining. The secondary anti-body mix was removed and larvae were again washed in PBS-T (6×10 min). Images were acquired using the Leica SP5 confocal microscope with a 20x water dipping lens (NA = 1.0) using a 405 nm and 633 nm laser.

### Neutral Red staining and image acquisition

Neutral red (NR) microglial staining was performed as previously described [186]. Briefly, live 3 or 5 dpf larvae were incubated in E3/PTU medium containing Neutral Red (NR) (Sigma-Aldrich) (2.5 µg/ml) for 2.5 h at 28 °C, and washed with E3/PTU for 30 min at 28 °C. Larvae were then anesthetized with 0.016% MS-222 and mounted dorsally in 1.8% low-melting-point agarose in E3. Z-plane images (2–4) were acquired using a Leica M165 FC microscope with a 10 × dry objective and a Leica DFC550 camera.

### Quantifications

All images were processed and quantified using the Fiji image processing package [335].

LysoTracker signal was quantified using thresholded maximum projections of the Z-stack. LT area and LT mean intensity (mean grey value) were calculated using the Measure tool. LT particles were analyzed using the Analyze Particle tool. Intracellular LT vesicles in microglia and oligodendrocytes were first selected within individual microglia or oligodendrocytes outlined by hand. LT+ particles within that outlined ROI were analyzed using a thresholded image with the Measure tool (to calculate %LT area in OLs). For analysis of LT+ vesicles within oligodendrocytes, 3-5 individual oligodendrocytes per fish were images and %LT area/OL was measured. The ten minute timelapse of LT+ vesicles in individual oligodendrocyte was quantified by manually counting the number of LT+ vesicles within oligodendrocytes (outlined by hand) every minute up until 10 minutes.

Myelin area was quantified using the Measure tool on thresholded maximum projections of the Z-stack in the hindbrain. Myelin sheaths of individual oligodendrocytes in the spinal cord were measured blindly using the Segmented Line tool or counted manually. For quantifications of *tmem106b;tfeb* fish, offspring required genotyping and therefore the genotype was unknown during analysis.

For NR+ microglia counting, serial images were stacked and NR+ microglia were counted by two independent researchers in the midbrain, since microglia first colonize the optic tectum in the midbrain around 2 dpf [46]. As NR staining was performed on progeny of heterozygous crosses, image acquisition and counting were performed blindly. For quantification of *mpeg*:GFP+ microglial morphology, maximum Z-projections of individual microglia were obtained (5 microglia/fish). Area and perimeter of microglia were measured on thresholded images using the Analyze Particles plug-in, and an average was calculated from 5 microglia/fish.

### Statistical testing

Statistical analysis was performed using PRISM GraphPad 8. Significance was calculated using Student’s t-tests, one-way ANOVA with Bonferroni multiple testing correction or two-way ANOVA with multiple testing correction. Data are presented as mean ± SD. A p value < 0.05 was considered significant.

## Supplementary Materials

**Suppl. Fig. 1.**
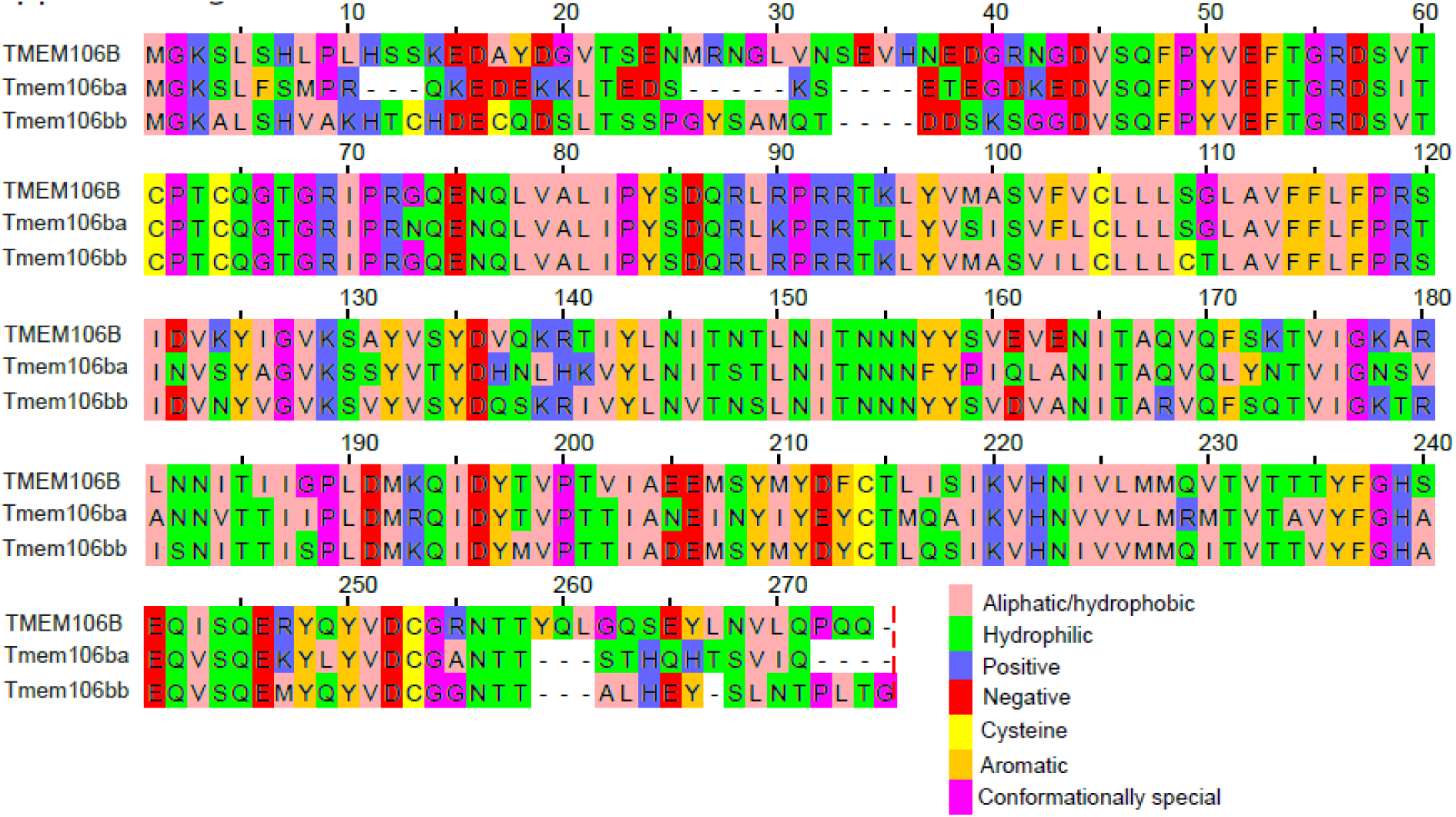
Zebrafish tmem106ba and tmem106bb are highly conserved compared to human TMEM106B protein. Conservation of human TMEM106B (top) compared to zebrafish Tmem106ba (middle) and Tmem106bb (bottom). Tmem106ba is 62% conserved, and Tmem106bb is 72% conserved compared to human TMEM106B. The residues are colored according to their physicochemical properties (see legend). Alignment and color code by JalView.

**Suppl. Fig. 2.**
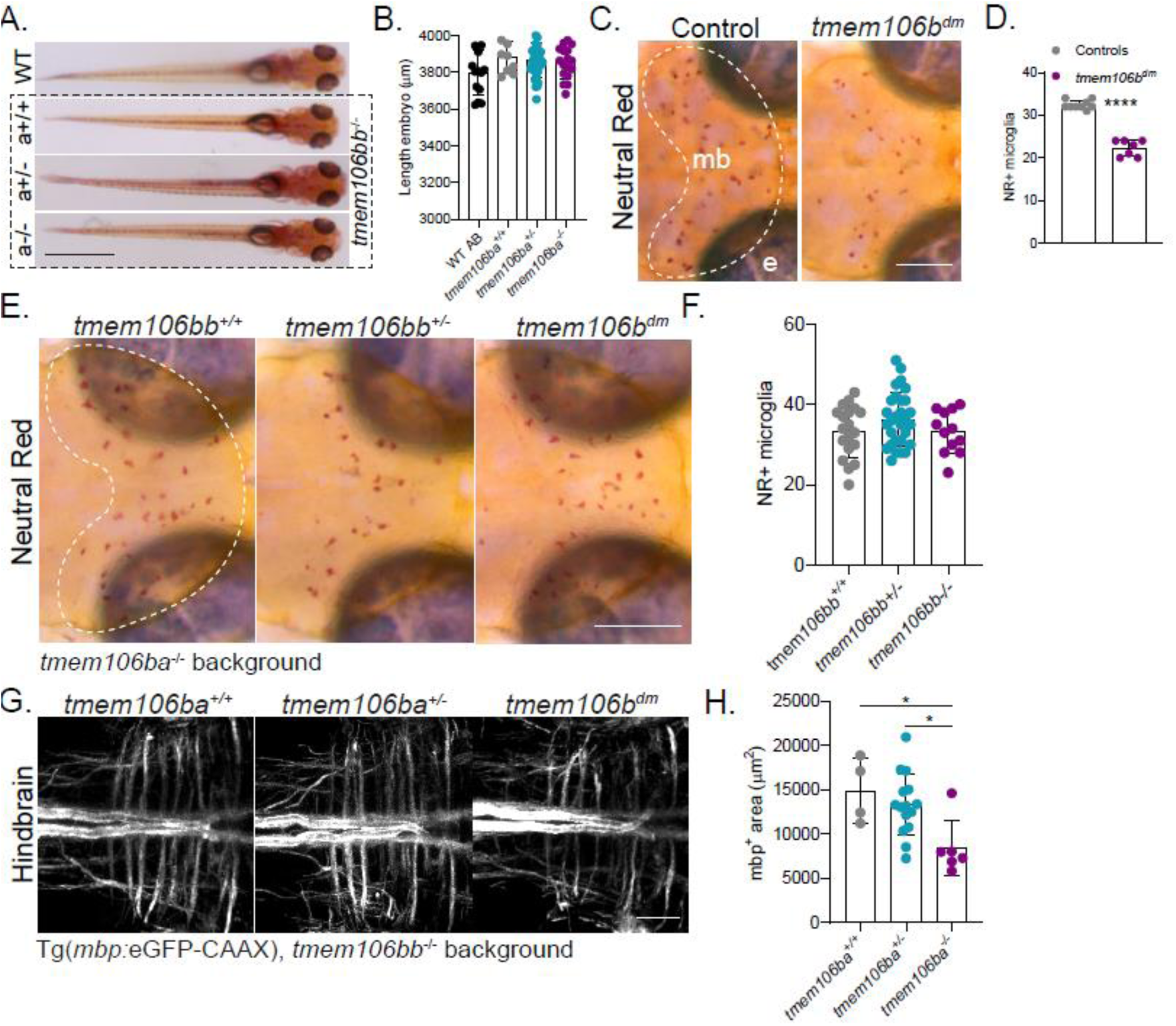
Supplemental data *tmem106b*^dm^ zebrafish. A) Dorsal representative images of WT AB and offspring of *tmem106ba^+/-^;bb^-/-^* incross of 5 dpf, showing no major developmental differences or delay. Scale bar equals 1000 µm. B) Quantification of the body length of 5 dpf larvae represented in A. C, D) Representative images of NR stained WT controls and *tmem106b^dm^* larvae at 3 dpf (C) and quantification of NR+ microglia in the midbrain (D). Scale bar equals 200 µm. E, F) Representative images of NR stained progeny of an incross of *tmem106ba^-/-^;bb^+/-^* at 3 dpf (E) and quantification of NR+ microglia in the midbrain (F). Scale bar equals 200 µm. G) Representative confocal images of mbp:GFP+ myelin in the hindbrain of progeny of a *tmem106ba^+/-^;bb^-/-^* incross. Scale bar equals 30 µm. H) Quantification of mbp:GFP+ myelin area in the hindbrain of larvae imaged in G. One-way ANOVA test with multiple comparisons or Student’s t-test was preformed to test for significance (p < 0.05). * p<0.05, **** p<0.0001.

**Suppl. Fig. 3.**
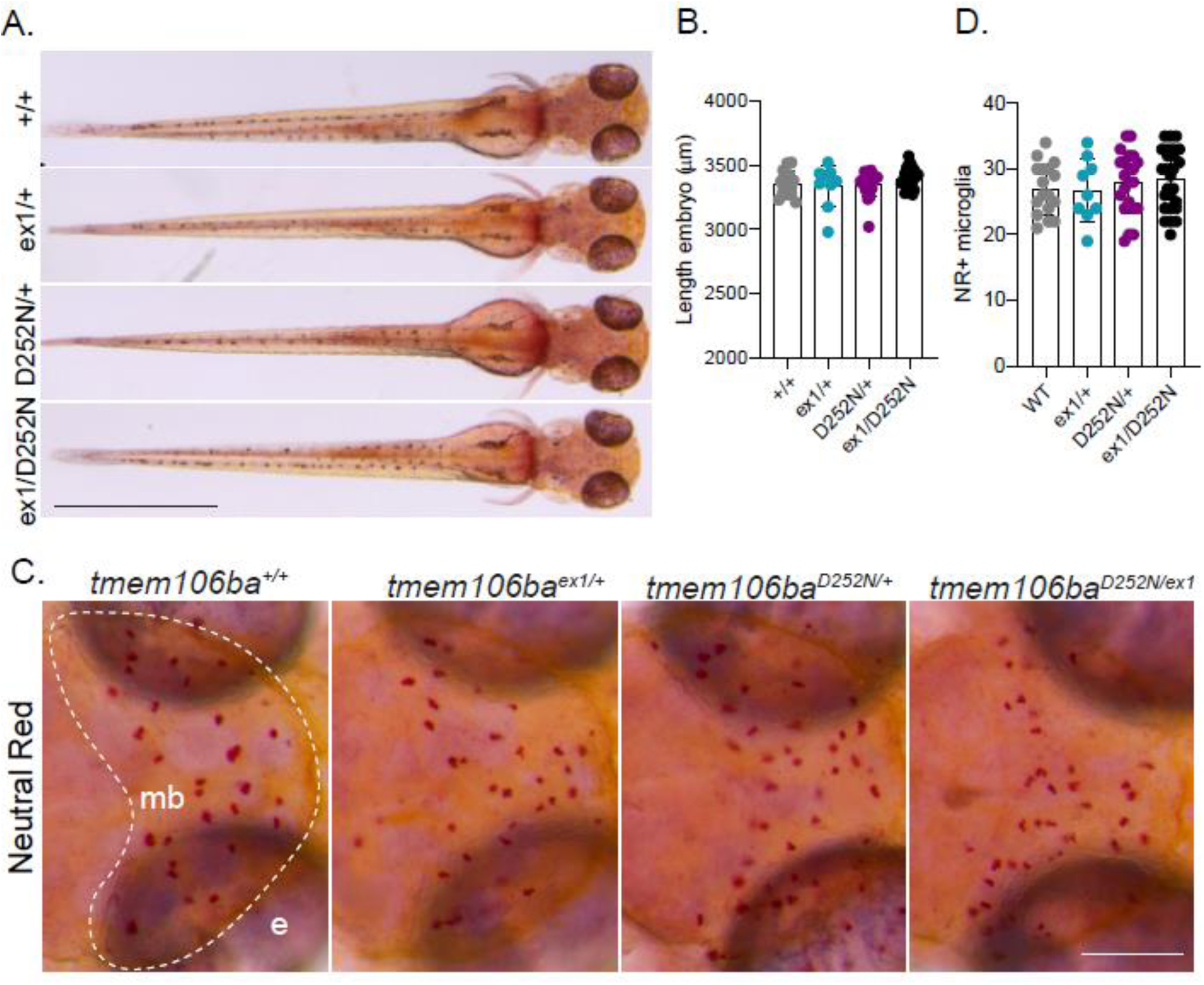
Supplemental data *tmem106b*^D252N/+^ zebrafish. A) Dorsal representative images of progeny of *tmem106ba^ex1/+^;b^-/-^* crossed with *tmem106ba^D252N/+^;b^-/-^* at 3 dpf, showing no major developmental differences or delay. B) Quantification of the body length of 3 dpf larvae represented in A. C, D) Representative images of NR stained progeny of *tmem106ba^ex1/+^;b^-/-^* crossed with *tmem106ba^D252N/+^;b^-/-^* at 3 dpf (C) and quantification of NR+ microglia in the midbrain (D). Scale bar equals 200 µm. One-way ANOVA test with multiple comparisons or Student’s t-test was preformed to test for significance (p < 0.05).

**Suppl. Fig. 4.**
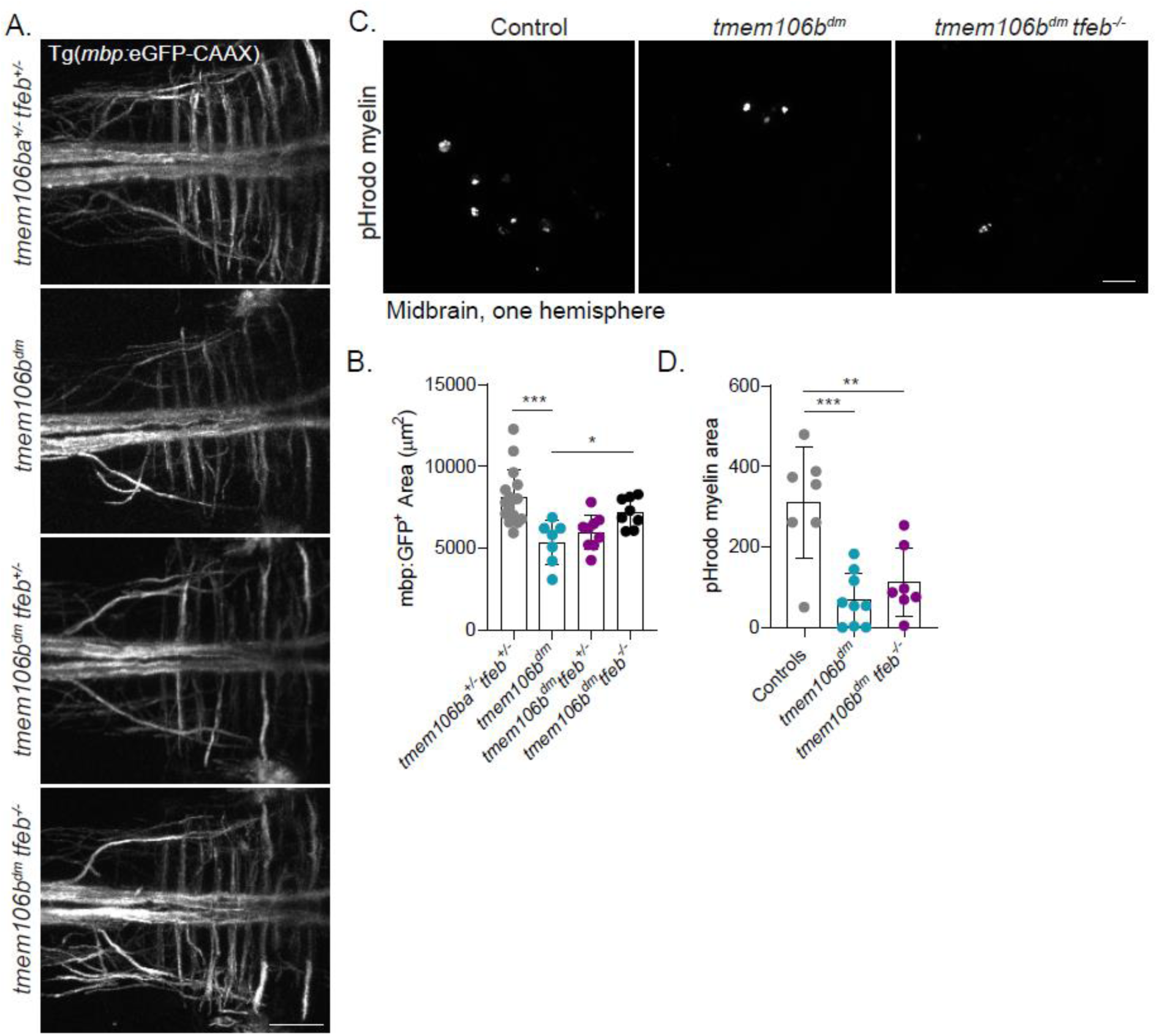
Supplemental data *tmem106bdm*;*tfeb* zebrafish. A, B) Representative in vivo confocal images (A) and quantification (B) of *mbp*:GFP+ myelin in the hindbrain of *tmem106ba*^+/-^ *tfeb*^+/-^ (n = 16), *tmem106b*^dm^ (n = 7), *tmem106b*^dm^ *tfeb*^+/-^ (n = 9) and *tmem106b*^dm^ *tfeb*^-/-^ (n = 8) larvae at 5 dpf. C, D) Representative in vivo confocal images (C) and quantification (D) of pHrodo-labelled myelin (magenta) in the midbrain of controls (n = 7), *tmem106b*^dm^ (n = 9) and *tmem106b*^dm^ *tfeb*^-/-^ (n = 7) larvae at 3 dpf after intracerebral injection (6 hpi). Scale bars equal 30 µm. One-way ANOVA test with multiple comparisons or Student’s t-test was preformed to test for significance (p < 0.05). * p<0.05, ** p<0.01, *** p<0.0001, **** p<0.0001.

## Acknowledgements

This work was supported by Metakids (NIW and TVH) and an Erasmus University Rotterdam fellowship (TVH). N.I.W. is a member of the European Reference Network for Rare Neurological Disorders (ERN-RND), project ID 739510. We thank the group of William Talbot (Stanford University) for *tfeb*-deficient and Tg(*cldnk*:Gal4/UAS:GFP-CAAX) zebrafish.

